# Estimates of quantal synaptic parameters in light of more complex vesicle pool models

**DOI:** 10.1101/2024.12.13.628305

**Authors:** Grit Bornschein, Simone Brachtendorf, Hartmut Schmidt

## Abstract

The subdivision of synaptic vesicles (SVs) into discrete pools is a leading concept of synaptic physiology. To better explain specific properties of transmission and plasticity, it has been suggested initially that the readily releasable pool (RRP) of SVs is subdivided into two parallel pools differing in their release probability. More recently, evidence was provided that sequential pools with a single RRP and a series-connected finite-size replacement pool (RP) inserted between the reserve pool (RSP) and RRP equally well or even better account for most aspects of transmission and plasticity. It was further suggest that a fraction of the presynaptic release sites (*N*) are initially unoccupied by SVs, with vesicle recruitment occurring rapidly during activity, and furthermore that the number of release sites itself changes with rapid dynamics during activity. Here we propose a framework that identifies specific signs of the presence of the series-connected RP, using a combination of two experimental electrophysiological standard methods, cumulative analysis (CumAna) and multiple probability fluctuation analysis (MPFA). In particular we show that if the y-intercept (y(0)) of CumAna is larger than *N* reported by MPFA (y(0) > *N*_MPFA_) this is a strong indication for a series-connected RP. This is due to the fact that y(0) reports the sum of RRP and RP. Our analysis further suggests that this result is not affected by unoccupied release sites, as such empty sites contribute to both estimates, y(0) and *N*_MPFA_. We discuss experimental findings and models in the recent literature in the light of our theoretical considerations.

## Introduction

The concept of Ca^2+^-dependent quantal vesicular release was introduced to synaptic physiology in the 1950s by Bernhard Katz and co-workers based on electrophysiological findings (Fatt and Katz, 1952;Del Castillo and Katz, 1954). The idea of quantal transmitter release received further support by the invention of rapid freezing combined with timed synaptic stimulation and electron microscopy (EM). Heuser and Reese succeeded in catching synaptic vesicles (SVs) in the act of fusion (Heuser et al., 1979). The concept of quantal release has since been confirmed in numerous studies and further extended (Neher, 2015). In this view, the amplitude of a postsynaptic current (PSC) is given by PSC = *p n q*, where *p* is the average probability that a vesicle (or “unit” as it was called initially) fuses, *n* is the number of fusion-competent SVs and *q* the amplitude of the postsynaptic response resulting from fusion of a single vesicle. The latter is referred to as the “quantal size” and the product of *p* and *n* is referred to as the “quantal content (QC)”. Hence, the amplitude of a PSC is the product of quantal content and quantal size and the overall rate of synaptic failures (F) or successes (S) strictly depends on the quantal parameters, being given by F = (1 - *p*)*^n^* and S = 1 – F.

The total number of fusion-competent vesicles constitutes the readily releasable pool (RRP) of SVs. Fusion occurs at specific presynaptic release sites (*N*) or docking sites (DS) as a synonym. However, quantitative estimates of RRP and *N* frequently deviate from each other and their relationship is not fully clear (Neher, 2015). The relationship between RRP and *N* and reasons underlying discrepancies between estimates of RRP and *N* are one of the topics of this article.

During synaptic activity, the RRP needs to be replenished by a continuous supply of SVs in order to maintain synaptic function. This supply comes from additional SV pools that have been postulated in recent years (Neher, 2015). SVs run through different Ca^2+^-dependent and - independent steps that include docking and priming to become fully fusion competent SVs of the RRP (Neher and Brose, 2018;Silva et al., 2021). Furthermore, the size of the RRP or the *N* can change during activity, which according to recent findings is a major determinant of synaptic short-term plasticity (STP; reviewed e.g. in (Neher and Brose, 2018;Miki, 2019;Schmidt, 2019;Neher, 2023)). Models with sequential pools of SVs have been proposed that accounted for several experimental findings on transmitter release and STP at different synapses in cerebellum (Miki et al., 2016;Doussau et al., 2017), neocortex (Bornschein et al., 2019b), brainstem (Lin et al., 2022) and hippocampus (Aldahabi et al., 2024).

We will distinguish three types of sequential models here. In the first type, presynaptic release sites are at rest fully occupied by SVs forming the RRP. During synaptic activity the RRP gets replenished from SVs that occupy replacement sites (RS) and form a replacement or replenishment pool (RP) with finite size that is intermediate between the reserve pool (RSP) and the RRP. The number of release sites itself can increase during high-frequency activity thereby giving rise to synaptic facilitation (Doussau et al., 2017). We will refer to this model as the sequential occupied sites (SOS RS/*N*) model. The second type is similar but at rest part of the release sites are considered to be not occupied by a SV. During high-frequency activity the occupancy but not the number of release sites increases thereby again accounting for synaptic facilitation (Miki et al., 2016). We will refer to this model as the sequential empty sites models here (SES RS/DS). The third model also has a fixed number of release sites and includes empty sites. In this model SVs are first in a loosely docked and primed state (LS) and reversibly pass through a tightly docked and primed state (TS) prior to fusion. An increase in TS accounts for facilitation here. This model is referred to as the LS/TS model (Neher and Brose, 2018;Neher, 2023). While SVs in the TS correspond to the RRP in the narrower sense, SVs in the LS are not identical to the RP vesicles (see Discussion).

Therefore, in order to account for short-term plasticity (STP), in these sequential models it is either assumed that the occupancy of a fixed number of *N* is incomplete at rest and increases during activity, i.e. the RRP increases but not the *N* (SES and LS/TS) (Trigo et al., 2012;Miki et al., 2016;Neher and Brose, 2018;Neher, 2023), or it is assumed that the number of *N* itself (= RRP) increases (SOS) (Valera et al., 2012;Brachtendorf et al., 2015;Doussau et al., 2017). In all models the increase in the size of the RRP is viewed as activity-dependent and reversibly. This is sometimes referred to as ‘overfilling’ of the RRP. In any case, the result is paired pulse facilitation (PPF) as observed e.g. at cerebellar parallel-fiber (PF) synapses. Results obtained with rapid freezing EM methods showed a reversible increase in the number of docked SVs shortly after timed synaptic stimulation (Kusick et al., 2020;Kusick et al., 2022), which is consistent with the sequential models and reversible overfilling. However, it does not differentiate between the three types of models.

An alternative and earlier model proposes a subdivision of the RRP into two parallel pools, differing in their vesicular release probabilities (*p*_v_) (Wölfel et al., 2007;Hallermann et al., 2010;Mahfooz et al., 2016). In traditional parallel models the intermediate RP is not implemented (Wölfel et al., 2007;Mahfooz et al., 2016). We refer to the traditional parallel model as PP model here. In recent work it was found that sequential and parallel models may almost equally well describe different aspects of experimental data (Eshra et al., 2021;Weichard et al., 2023). Therefore, it is not only difficult to distinguish between the different sequential models but also between sequential and parallel models.

Two standard electrophysiological methods are frequently used for the quantitative estimation of synaptic parameters, including SV pools: The analysis of cumulative PSC amplitude plots (CumAna) (Schneggenburger et al., 1999) and multiple probability fluctuation analysis (MPFA) (Clements and Silver, 2000). In CumAna trains of action potentials (APs) are applied under recording conditions suitable to drive synapses into an equilibrium between release and replenishment. This is typically given, if the vesicular release probability (*p*_v_) is sufficiently high to achieve a steady-state depression of ∼60% during the train. At synapses with moderate *p*_v_ this might require to perform recordings in an elevated extracellular Ca^2+^ concentration ([Ca^2+^]_e_). Additional methodological requirements for successful application of CumAna have been described in detail elsewhere (Neher, 2015). The result is a linear relationship between cumulative PSC amplitudes late in the train and the corresponding stimulus numbers or times. Fitting a line to this linear phase and extrapolating it to the y-intercept (y(0)) removes the contribution of SVs added via replenishment in the steady-state phase. Hence, the y-intercept is thought to report the initial size of the RRP. More precisely, it reports a value close to the decrement of the RRP during the train and the “real” size of the RRP can be obtained with a correction calculation as long as the RRP gets strongly depleted during the train (Thanawala and Regehr, 2013;Neher, 2015). Dividing the first PSC or QC by the y-intercept then gives the average *p*_v_.

For MPFA, PSCs are recorded under conditions of several different *p* values, which is typically obtained by changing the [Ca^2+^]_e_. Alternative approaches to change *p* include broadening the presynaptic AP, e.g. by application of blockers of voltage gated potassium channels like TEA or 4-AP. The variance and the mean of the PSCs are calculated during stable recording periods after wash-in of each different Ca^2+^ solution or alternative treatments and the variance is plotted against the mean. A parabolic fit to these data yields *N* and the average release probability per release site (*p*_N_) (Clements and Silver, 2000).

In our experiments at cortical synapses we typically found that the estimates of RRP from the y-intercept were larger than the estimates of *N* from MPFA for the same synapses. Accordingly, the estimates of the *p*-values showed the opposite behavior (Schmidt et al., 2013;Baur et al., 2015;Bornschein et al., 2019a;Bornschein et al., 2019b). Deviations between y-intercept and *N* were observed also at many other synapses (Neher, 2015). However, the cause of these deviations is not fully clear.

CumAna and MPFA were established before the development of sequential or parallel pool models. Here, we used computer simulations to systematically investigate which entities are actually reported by the two experimental methods in light of different pool models and considering incompletely populated release sites. The systematic approach starts with basic simulations of simple arrangements of SV pools, which become increasingly complex. This article aims to investigate how much information about the organization of SVs can be obtained based on the experimental results alone, without the need to fit complex models to the data. We suggest that a combination of CumAna with MPFA provides complementary insights into the functional organization of SV pools and their dynamics that cannot be achieved with either method alone.

## Methods

### Computer simulations

All simulations were performed in Mathematica 14 (Wolfram) as described in more detail elsewhere (Wender et al., 2023).

#### Algebraic simulations

For the algebraic simulations of CumAna it was assumed that the vesicular release probability *p*_v_ remains constant during a train of APs. The number of release sites occupied by a releasable vesicle (*n*[*i*]) during the i^th^ pulse of a train was calculated as

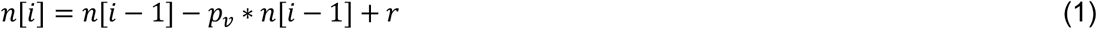

where *p*_v_ \**n*[*i-1*] is the release in the preceding pulse and *r* is the recruitment rate from the RSP. Quantal content of the i^th^ pulse is given by *n*[*i*] * *p*_v_. In simulations mimicking the presence of a series-connected RP, the RP was simulated as

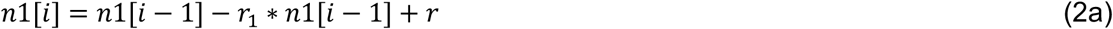

and the RRP by

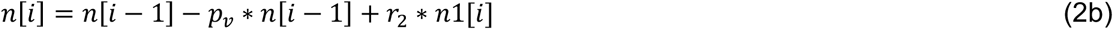

Two parallel pools were simulated using equation (1) for each RRP and the *n*[*i*] from the two pools were summed linearly, assuming independence of release sites.

In the presence of replenishment, the y-intercept (y(0)) underestimates the size of the RRP but can be corrected, using the following formula (Neher, 2015):

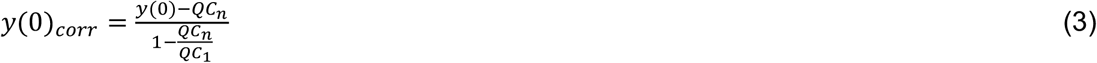

where QC are the quantal contents during the last (n) or first stimulation, respectively. An alternative correction (Thanawala and Regehr, 2013) assumes a restricted number of release sites, while our simulations allowed for variability in the *N* or their occupancy according to recent models of STP (Miki et al., 2016;Doussau et al., 2017;Bornschein et al., 2019b;Lin et al., 2022;Aldahabi et al., 2024). Hence, the alternative correction was not considered further here.

#### Stochastic simulations

For Monte Carlo simulations of release for MPFA and CumAna, random real numbers were generated using ‘RandomReal’ of Mathematica. Release sites were simulated individually. Release from a given release site occurred only if the conditions set by *p*_v_ or *p*_N_ and *p*_occ_ were met. If a release site had released, release from this site could occur again, only if the conditions of *p*_v_ or *p*_N_ and *p*_repl_ were met. To give the total response, quantal contents were assumed to add linearly across release sites.

## Results

### Basic simulations of CumAna and MPFA

For CumAna, we started with a simple algebraic simulation of a single RRP set to 10 SVs with *p*_v_ set to 0.6 in the absence of any replenishment (**Fig. 1A**). All release sites were assumed to be fully occupied, i.e. the RRP corresponded to *N* in these simulations. During a train of APs, the RRP was rapidly exhausted, giving rise to a horizontal line fit in the CumAna plot. In this most simple scenario, the y-intercept correctly reports the RRP size of 10 and dividing the first QC by the y-intercept (QC1/y(0)) gives the set value of *p*_v_ of 0.6. Hence, in this very simple scenario the nominal values and the actual values reported by CumAna perfectly match. The slope of the line is zero, according to the absence of replenishment.

**Figure 1.**
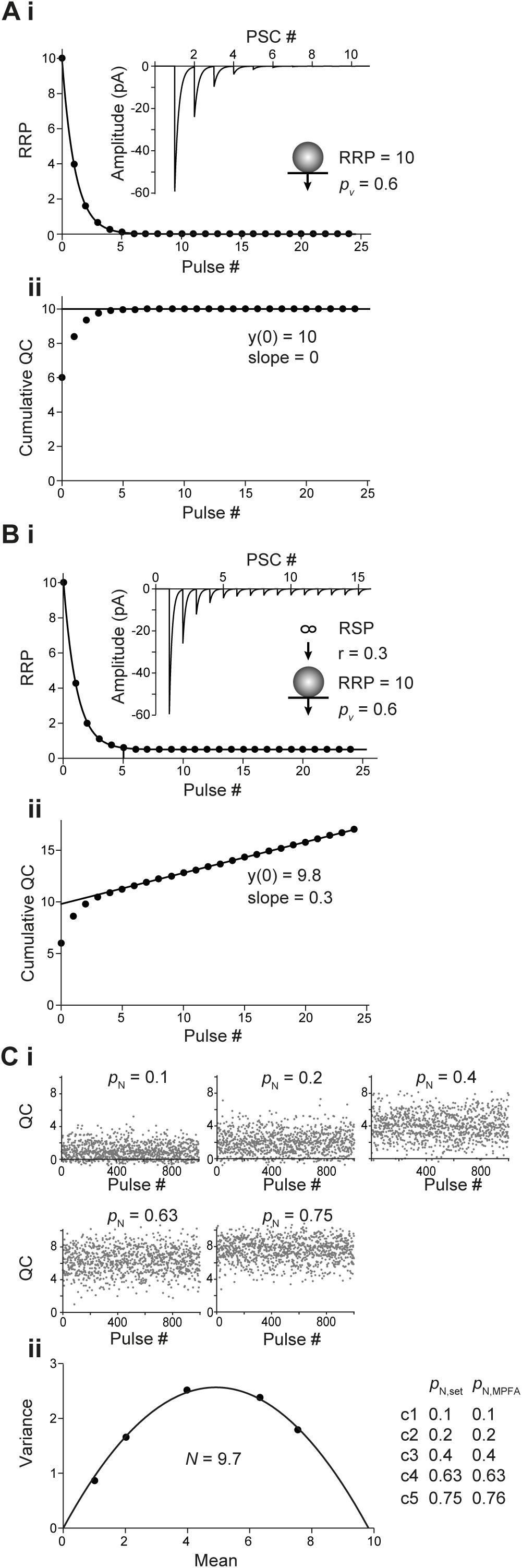
Basic analysis with CumAna and MPFA. **Ai.** Simulation with completely occupied release sites, a single RRP of 10 SVs (*top*) in the absence of replenishment and *p*_v_ of 0.6 remaining constant during the train. The RRP rapidly dropped to zero due to the absence of replenishment during a train of 25 APs. *Inset*: Simulated PSCs assuming a *q* of 10 pA. Scheme illustrating vesicular release from the RRP with *p*_v_. **ii.** Cumulative quantal contents were plotted against pulse number starting at 0 and a line was fitted to the last 5 data points and back-extrapolated to the y-axis. Note that the y-intercept (y(0)) correctly reports the RRP size of 10 SVs. **Bi.** As in (**A**) but in this simulation replenishment of the RRP occurred from an infinite RSP with rate constant r of 0.3 (top). **ii.** CumAna as above. The y-intercept is 9.8 and the slope of the line is 0.3. **Ci.** For MPFA 1000 release processes were simulated stochastically at five different settings for *p*_N_ as indicated with *N* set to 10. Quantal contents are shown plotted against the pulse number. Note that the variance of the quantal contents first increases and then decreases whereas the mean content constantly increases as a function of *p*_N_. **ii.** Variances were plotted against the corresponding mean quantal contents and fitted by a parabola. The *N* estimated by the parabolic fit was close to the set value and the set values for *p*_N_ were correctly reported by the MPFA.

Next, the simulation was extended to include constant replenishment with rate constant r of 0.3 SVs/stimulus from an infinite RSP (**Fig. 1B**). In this scenario, the slope of the line fit reported the r of 0.3. However, the y-intercept now slightly underestimated the size of the RRP, being 9.8 rather than 10. Accordingly, *p*_v_ was slightly overestimated. For this scenario, the y-intercept can be corrected by analytical calculations (**formula 3** in Methods) to yield the set value of the RRP of 10 and to derive the correct *p*_v_ of 0.6 (Neher, 2015).

For MPFA, we started with stochastic simulations of 10 fully occupied *N* with homogenous release probability (*p*_N_), which were assumed to be fully replenished between pulses. (**Fig. 1C**). The *p*_N_ was set to different values such that the parabola was well defined, i.e. the larger *p*_N_ exceeded 0.5, which is the apex of the parabola that has to be passed for a reliable fit (Clements and Silver, 2000). The parabolic fit resulted in an *N*_MPFA_ of ∼10, which is the intercept of the parabola with the x-axis. Depending on the number of simulated release processes, the x-axis intercept fluctuated more or less strongly around the set value of 10. Even with the 1,000 runs used in these simulations, which cannot be achieved experimentally, the deviation from the set value was up to 0.3 (**Fig. 1C**). The deviation from the set value was reduced to 0.01 after 10,000 runs. The *p*_N_ obtained from the parabolic fits reliably reported the set values even with 1,000 runs. In summary, in these basic scenarios, both CumAna and MPFA provide reliable estimates of the “real values” for RRP, number of release sites and *p* that are congruent between the two analysis methods.

### CumAna in SOS and PP model

To investigate CumAna for the SOS model, we inserted the finite-sized RP between RSP and RRP in the simulation (**Fig. 2**). As above, all release sites were assumed to be fully occupied initially (Doussau et al., 2017). The *p*_v_ of SVs in the RRP was set to 0.6 as above. The sum of SVs in RP and RRP was set to 10. The transition of SVs from the infinite RSP to the finite RP occurred with rate constant r_2_. The transition of SVs from RP to RRP occurred with a faster rate constant r_1_. r_1_ was either only moderately faster than r_2_, which resulted in immediate depression (**Fig. 2A**), or it was much faster than r_2_, leading to overfilling of the RRP during the first APs and initial facilitation despite the high *p*_v_ (**Fig. 2B**). These models thus simulate experimental results and their interpretation, where a key determinant of STP is the transition rate of SVs from a series-connected RP to the RRP (Miki et al., 2016;Doussau et al., 2017;Schmidt, 2019).

**Figure 2.**
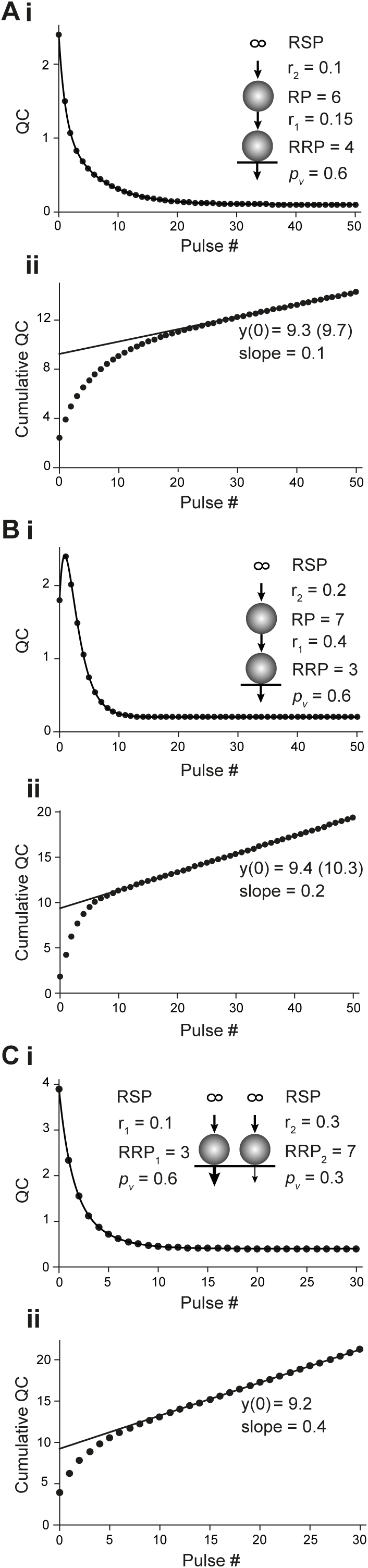
CumAna in the SOS versus PP model. **Ai.** Plot of the quantal content (QC) against pulse number fitted with a double exponential function. SOS model of a depressing synapse. The size of RRP and RP were 4 and 6, respectively. Replenishment from the infinite RSP to RP occurred with r_2_ of 0.1 and the transition of SVs from RP to RRP with r_1_ of 0.15. *p*_v_ was set to 0.6 throughout the train. *Inset*: Scheme illustrating vesicular release and replenishment of SVs in the SOS model. **ii.** Cumulative quantal content plotted as a function of pulse number. The back-extrapolated line fit to the last 5 data points has a y-intercept at 9.3 (9.7 with correction according to Neher (2015)). Note that this is close to the sum of RP and RRP. Accordingly, dividing the first quantal content by the y-intercept gives a value of 0.26 (0.25) that underestimates *p*_v_. The slope of the line fit is 0.1, i.e. it corresponds to r_2_. **Bi.** As in (**Ai**) but for a facilitating synapse. The set values in this simulation were as follows: RRP = 3; RP = 7; r_1_ = 0.4; r_2_ = 0.2; *p*_v_ = 0.6. Note the initial facilitation despite the high *p*_v_, which is due to an increase in the RRP (overfilling). **ii.** Cumulative quantal content as function of pulse number. The line fit has a y-intercept of 9.4 (10.3 following correction) and a slope equal to r_2_. Hence, the y-intercept is again close to RP plus RRP and the quantity of first quantal content / y-intercept yields 0.19 (0.17), again underestimating *p*_v_. **Ci.** As in (**Ai**) but for the PP model of a depressing synapse. RRP_1_ and RRP_2_ harbored 3 and 7 SVs with *p*_v_ of 0.6 (thick arrow) and 0.3 (thin arrow), respectively. RRP_1_ was replenished slower with r_1_ = 0.1 and RRP_2_ faster with r_2_ = 0.3 SVs/pulse. *Inset*: Scheme illustrating vesicular release and replenishment of SVs in the parallel model. **ii.** As in (**Aii**). y-intercept = 9.2 (9.8 following correction), slope = 0.4 (corresponding to r_1_ + r_2_), calculated release probability = 0.42 (0.4).

The curve of the decrease in QC in these simulations was biphasic (**Fig. 2Ai,Bi**), which is due to the presence of the RP with finite size. RRP will show the same behavior, but scaled by *p*_v_. This is a clear deviation from the monophasic decrease in QC observed in the simulations in which the RRP was directly replenished from the RSP (**Fig. 1**). A biexponential decay can thus indicate the presence of a finite-sized RP (Bornschein et al., 2019a) but it could also indicate a subdivision of the RRP into two parallel pools (see below).

We found that the fitting line of the corresponding CumAna plots had a slope of r_2_. The slope was independent of r_1_ and solely determined by the rate-limiting transition rate r_2_. Remarkably, the y-intercept did not correspond to the size of the RRP, but had a value close to 10. Thus, the y-intercept actually reflects a value close to the sum of RP and RRP. Accordingly, the *p*_v_ value calculated from the ratio of QC1 or PSC1 to the y-intercept significantly underestimated the specified value of 0.6 (**Fig. 2Aii, Bii**).

For CumAna with the PP model, the RRP was subdivided into two subpools with different *p*_v_ and replenishment rates (Wölfel et al., 2007;Mahfooz et al., 2016). RRP1 harbored 3 SVs with high *p*_v1_ of 0.6 and slower r_1_ of 0.1 SVs/stimulus, while RRP 2 harbored 7 SVs with lower *p*_v2_ of 0.3 but faster replenishment with rate constant r_2_ of 0.3 (**Fig. 2C**). As for the above sequential model, the time course of the decrease in quantal content was biexponential (**Fig. 2Ci**). The line fit in the cumulative quantal content plot had a slope of 0.4, i.e. a value corresponding to the linear sum of r_1_ and r_2_. The y-intercept had a value close to the total RRP, which means that a larger part of the pool with a low *p*_v_ value showed up in the y-intercept (**Fig. 2Cii**). Consequently, the *p*_v_ as reported by dividing the first quantal content by the y-intercept was close to the average between *p*_v1_ and *p*_v2_.

Taken together, these results indicate that biexponential decays in QC or PSC amplitudes during train experiments hint towards the presence of a more complex organization of pools of SVs as proposed in the sequential or the parallel model. However, CumAna alone cannot identify these pools since there is no a priori knowledge about the presence of a RP or a subdivision of the RRP. Moreover, the sequential and the parallel model lead to similar results in CumAna. In particular, for the SOS model the y-intercept reports the sum of RRP and RP (y(0) = RRP + RP) and in the PP model it reports the sum of the high and low *p*_v_ pools (y(0)= RRP_high *p*_ + RRP_low *p*_).

### CumAna in empty sites models

In the next step, we used stochastic simulations to explore the outcome of CumAna if the initial occupancy of the release sites is incomplete (Miki et al., 2016), which we refer to as empty sites models (ES). First, we simulated a single RRP model with a fractional release site occupancy of 0.7 in the absence of replenishment (**Fig. 3**). *N* was set to 10 with a corresponding RRP of 7 and *p*_v_ was set again to 0.6 (**Fig. 1**). Interestingly, the y-intercept of CumAna reported the value of 10 in this scenario. Thus, although the RRP harbored only 7 SVs on average in these simulations, the y-intercept reports the set value for *N* of 10 (**Fig. 3A**). However, due to the reduced occupancy and the correspondingly smaller RRP the quantal content of the first PSC was only 4.4 rather than 6. Accordingly, the ratio of QC1/y(0) underestimated the true *p*_v_ and was only 0.44 instead of 0.6. This value is close to the set value of *p*_v_ of 0.6 multiplied by the occupancy of 0.7. Hence, in ES models *p*_v_ does not report the intrinsic fusion probability of a SV but rather the fusion probability multiplied by the occupancy (cf. Neher, 2023).

**Figure 3.**
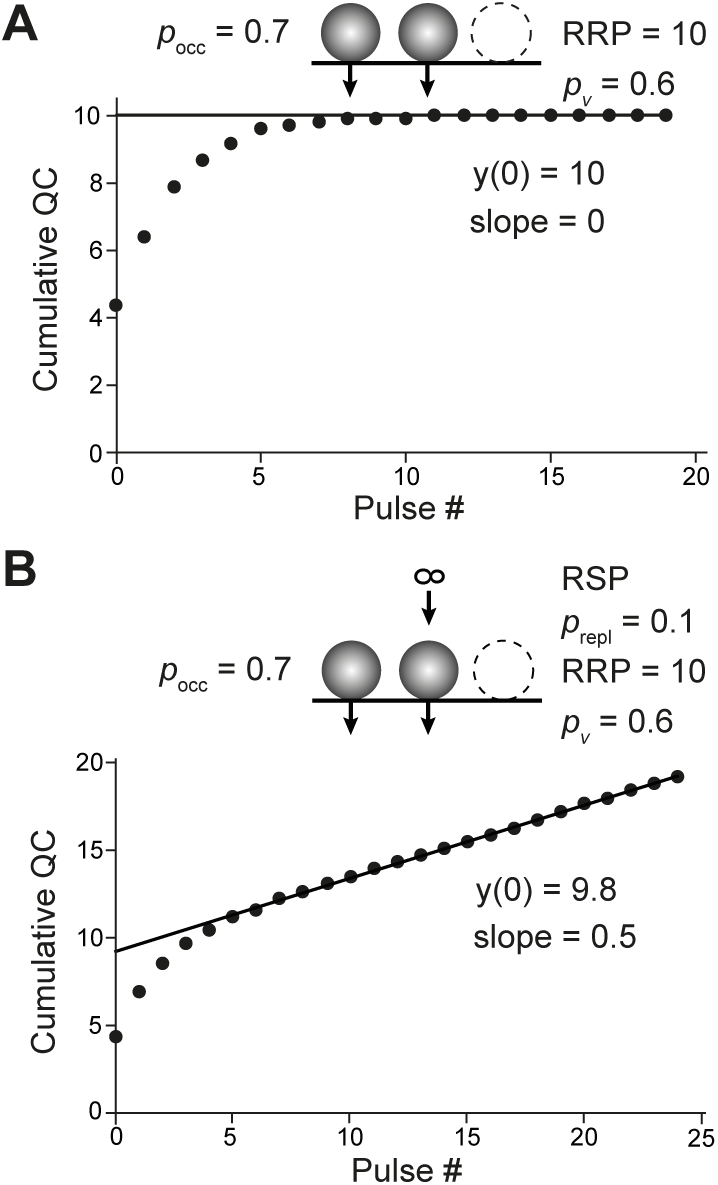
CumAna with incomplete initial occupancy of release sites. **A.** Stochastic simulation of quantal contents in the absence of replenishment during a train of 20 APs. The train was repeated 10 times and the average quantal content plotted as a function of pulse number. Further settings were as follows: *N* = 10; initial *p*_occ_ = 0.7; *p*_v_ = 0.6, assumed to be constant during the train. The line fit yielded a y-intercept of 10, thus, a value reflecting the total number of release sites rather than the RRP of 7. *Inset*: Scheme illustrating the RRP with *p*_occ_ and the vesicular release from the RRP with *p*_v_. **B.** As in (**A**) but for 100 APs (only the first 25 are shown) with replenishment from an infinite RSP (scheme in the *inset*). A release site that had released in a preceding pulse was replenished with probability *p*_repl_ = 0.1. Between pulses one and two the fractional occupancy of the release sites was assumed to increase to one.

Next, the ES simulation was extended by a single replenishment step from the RSP (**Fig. 3B**). In this scenario, the y-intercept reported a value of 9.8, i.e. a value close to but somewhat smaller than the set value for the number of release sites. Accordingly, the ratio of QC1/y(0) again underestimated the true *p*_v_ and was 0.43. Depending on the details of the settings for *p*_v_, initial occupancy and increase in occupancy between pulses, this model could either produce initial facilitation followed by depression or immediate depression (not shown). Yet, the principle result that the y-intercept reports a value close to the total number of release sites, including those that were empty initially, rather than the RRP was not affected by these differences. Finally, the ES simulation was extended by the intermediate RP with finite size, resulting in the SES model. As expected from the previous section, the y-intercept now includes the RP, i.e. in the SES model y(0) reports the sum of RP, RRP and empty sites (**Table 1**). Accordingly, *p*_v_, calculated from QC1 and y(0), will be smaller than the product of fusion probability and release site occupancy.

**Table 1.**
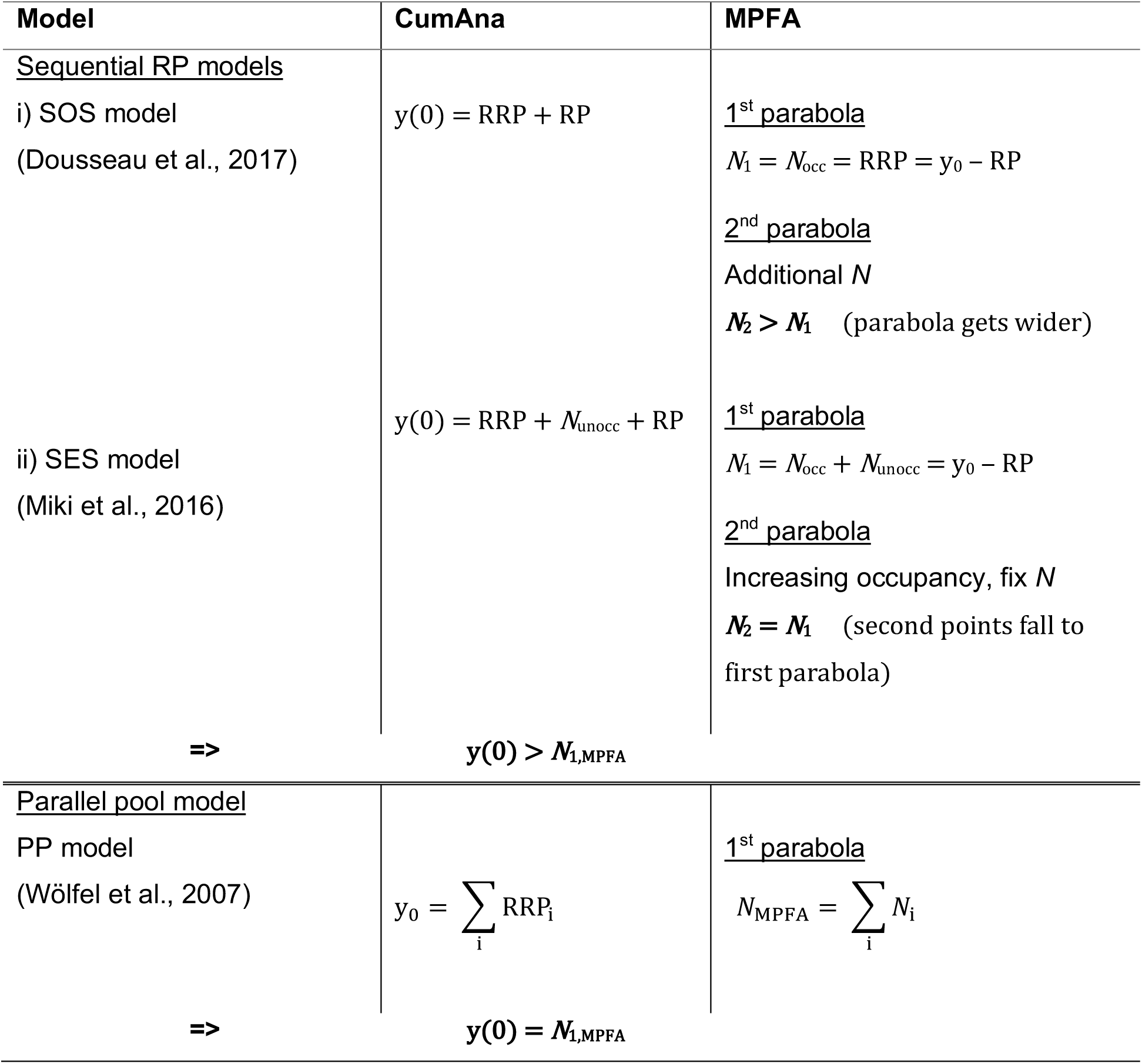
Summary of simulation results for SOS, SES and PP models. The LS/TS model was not covered in the simulations. With regard to y(0) and *N*_1,MPFA_ the following results are expected (Neher and Brose, 2018;Neher, 2023): y(0) = LS + TS + *N*_unocc_ and *N*_1,MPFA_ = *N*_LS_ + *N*_TS_ + *N*_unocc_, i.e. y(0) = *N*_1,MPFA_ as in the PP model (see **Fig. 6**).

Taken together, if the initial occupancy of the release sites is incomplete (ES models), the y-intercept reports the total *N* including empty sites rather than the RRP (y(0) ≈ *N* > RRP) and the quantity QC1/y(0) underestimates *p*_v_. In fact, the value of *p*_v_ from the CumAna corresponds to the product of the set value of *p*_v_ (intrinsic vesicular fusion probability) and the occupancy. However, for the SES model the RP will add to y(0) such that y(0) ≈ *N* + RP and the reported *p*_v_ will be smaller than fusion probability times occupancy. It should be noted at this point that RP contributes to y(0) in both sequential models, SOS and SES. We further have to note that there is no a priori knowledge of the actual occupancy of the release sites and therefore CumAna alone cannot identify incomplete occupation of release sites. In the following sections we therefore investigated whether MPFA could be useful to help distinguishing increasing release site occupancy as mechanism of facilitation from a factual increase in the number of release sites.

### MPFA with paired pulses in SOS and PP model

As indicated by the above results CumAna alone cannot distinguish between the different models. Hence, we proceeded by simulating MPFA to see whether a combination of CumAna and MPFA will yield deeper insights. First, we used the SOS model and simulated MPFA with paired pulses at a short interstimulus interval (ISI) and a longer interval between the double pulses that allowed the synapse to return to its initial state, i.e. complete relaxation from any STP during the paired pulses with a complete recovery of the RRP to its initial size. We simulated the following two scenarios: First, full initial occupancy of release sites with an actual increase or decrease in the number of *N* in the second pulse (**Fig. 4A-C**). Second, full initial occupancy of release sites with increasing or decreasing *p*_N_ in the second pulse, while *N* stayed constant (**Fig. 4D-E**).

**Figure 4.**
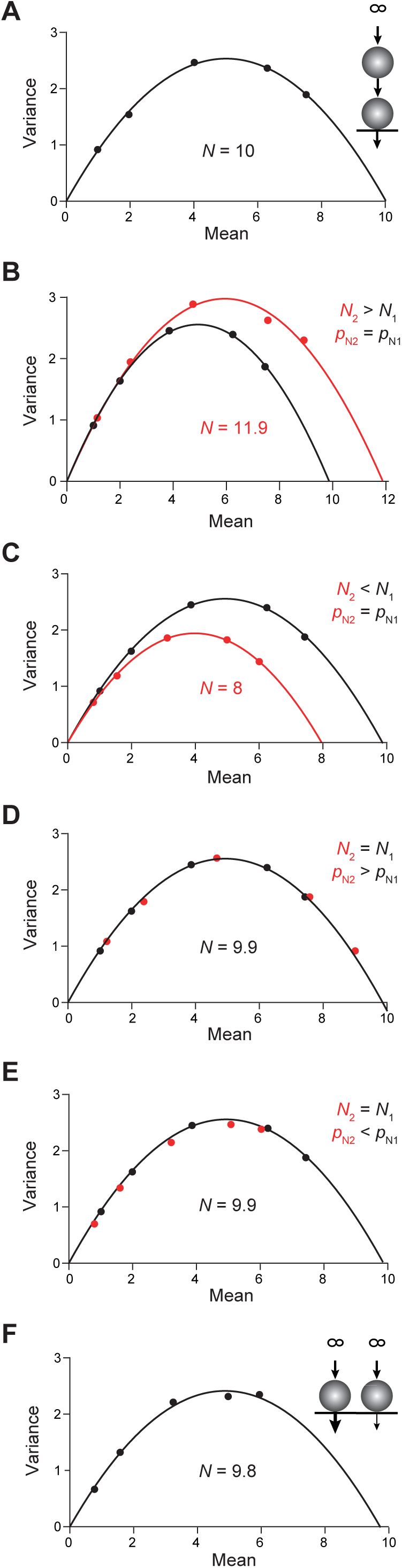
MPFA in the sequential versus parallel model. **A.** Sequential model with an RP and STP resulting from changes in *N* or *p*_N_. MPFA of the first pulse as described in **Fig. 1C**. It is assumed that SVs from the RP do not contribute in the first release process. The parabolic fit yielded an *N* very close to the set value of 10. *Inset*: Scheme illustrating vesicular release and replenishment of SVs in the sequential model. **B.** Black parabola and simulations as in (**A**). The red parabola is the result for *N* being increased to a value of 12 in the second pulse due to rapid recruitment of new *N* from the RP. The *p*_N_ values were as in (**A**). **C.** As in (**B**) but for *N* being reduced to 8 in the second pulse due to slow replenishment. **D.** Black points and parabola as in (**A**). The red points are the result of *p*_N1_ being increased in second pulse (*p*_N2_ : 0.12, 0.24, 0.48, 0.76, 0.9), while *N* was kept constant. Note that the red points fall to the initial black parabola albeit with a rightward shift i.e. towards higher release probabilities. **E.** Same as in (**D**) but for reduced *p*_N_ in the second pulse (0.08, 0.16, 0.32, 0.5, 0.6). **F.** PP model for two RRPs occupying release sites *N*_1_ (3) and *N*_2_ (7) with *p*_N1_ (thick arrow) as in (**A**) and *p*_N2_ = 0.7 * *p*_N1_ (thin arrow). MPFA as described in **Fig. 1C**. The parabolic fit yielded *N* of 9.8, which is close to the set value of *N*_1_ + *N*_2_. . *Inset*: Scheme illustrating vesicular release and replenishment of SVs in the parallel model.

In the first scenario, a parabola fitted to the mean – variance plot of the first PSC amplitudes or quantal contents yielded the nominal value for the number of release sites *N* as in the above simulations (**Figs. 1C, 4A**). This value of *N* corresponds to the size of the RRP and excludes the RP. Thus, in comparison to CumAna with the SOS model the *N* yielded by the first parabola of MPFA is smaller than the y-intercept. The second parabola deviated from the first one in characteristic ways: If *N* increased between paired pulses, the parabola got wider, whereas it got narrower if *N* decreased (**Fig. 4B, C**). By contrast, in the second scenario with changes in *p*_N_ rather than in *N*, the points of the second pulses fell to the first parabola. If *p*_N_ was increased they were shifted towards higher mean values and if *p*_N_ was decreased they shifted towards smaller means (**Fig. 4D, E**). The results for the second parabolas are consistent with previously published results (Clements and Silver, 2000).

Next, we performed the MPFA with the PP model (**Fig. 4F**). A total *N* of 10 was subdivided into *N*_1_ and *N*_2_. *N*_1_ had a size of 3 with *p*_N1_ set to 0.1, 0.2, 0.4, 0.63 or 0.75 to mimic e.g. wash-in of increasing [Ca^2+^]_e_. *N*_2_ was set to 7 with *p*_N2_ set to 0.7 * *p*_N1_, representing the pool with lower release probability. In this case *N*_MPFA_ reported by the parabolic fit was close to 10, i.e. it reported both, the high and the low *p*_N_ sites. Thus, for parallel pools the y-intercept obtained by CumAna and the *N*_MPFA_ derived by MPFA will be very similar, being identical in theory.

Taken together, these simulations indicate that a combination of CumAna and MPFA can provide a means to distinguish between SOS and PP models. In particular, for SOS the y-intercept will be larger than *N* estimated by MPFA (y(0) > *N_MPFA_*), whereas for PP the y-intercept and *N* will be equal (y(0) = *N_MPFA_*; **Table 1**).

### MPFA with paired pulses in the ES models

We proceeded by analyzing what happens in MPFA if the source of STP is a change in the occupancy of release sites rather than a change in the number of release sites. In order to analyze such incomplete occupancy of release sites, we started with paired pulses in a presynaptic depletion model in which all *N* were initially fully occupied but emptied sites were not replenished prior to the second pulse, i.e. the *N* were more or less depleted in the second pulse, depending on the quantal content released in the first pulse (**Fig. 5A**). The total *N* in the simulation remained constant, i.e. merely the occupancy of release sites decreased between pulses one and two. In this scenario all points of the second pulse fell to the first parabola albeit at apparently lower *p*_N_ values. The shift of the second points was strongest for the highest nominal *p*_N_ settings although the set values for *p*_N_ were not changed between pulses, i.e. with regard to the nominal values *p*_N2_ was equal to *p*_N1_. In fact, the actual values for *p*_N2_ were equal to the set values for *p*_N2_ multiplied by the actual occupancy of the *N* in the second pulse (**Fig. 5A**).

**Figure 5.**
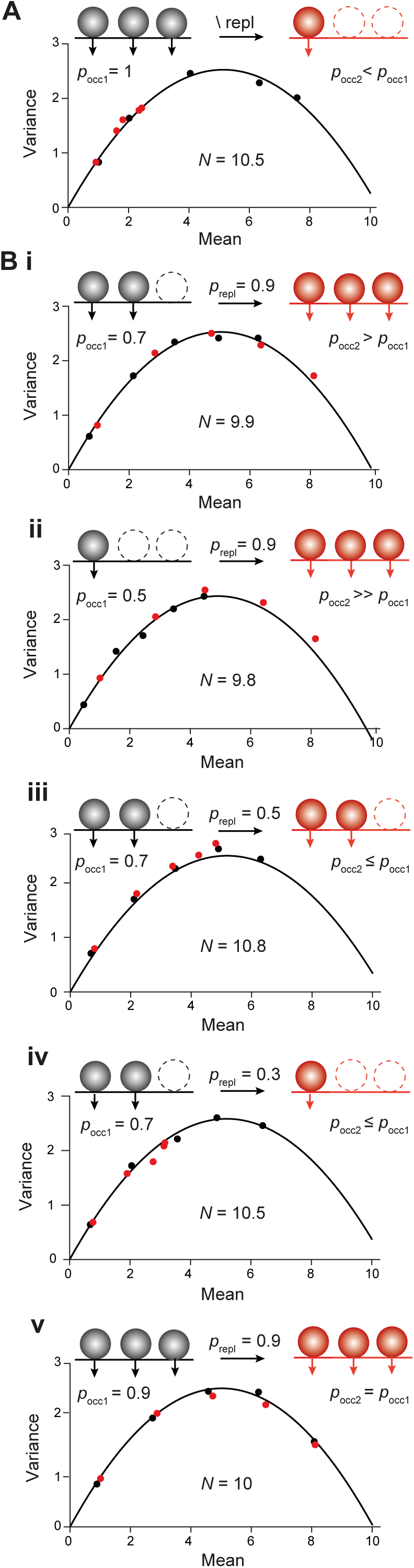
MPFA with STP resulting from changes in the occupancy of *N*. **A.** MPFA as in **Fig. 1C**, **4A** but for paired pulses. In the first pulse the *N* were fully occupied by SVs (black points and parabola). Release sites were not replenished in the second pulse, resulting in presynaptic depletion of SVs (red points). Note that the red points fall to the initial parabola albeit at apparently lower *p*_N_ values. *Inset*: Scheme illustrating occupancy of *N* by SVs in the first (*p*_occ1_, black spheres) and the second pulse (*p*_occ2_, red spheres) without replenishment (\ repl) in between. Empty release sites are shown with dashed lines. **B.** MPFA with paired pulses as in (**A**) but for different probabilities of initial occupancy of *N* (*p*_occ1_) with different probabilities of replenishment between pulses (*p*_repl_). *Inset* **i-v**: Scheme illustrating occupancy of *N* by SVs in the first (black circles) and the second pulse (*p*_occ2_, red circles) with replenishment between pulses**i.** *p*_occ1_ of 0.7 and *p*_repl_ of 0.9. The first parabola (black) estimates *N* according to the set value of 10. The points of the first (black) and the second pulse (red) fall to the same parabola at apparent *p*_N_ values that are given by the nominal *p*_N_ multiplied with the occupancy in the corresponding pulse. **ii.** *p*_occ1_ of 0.5 and *p*_repl_ of 0.9. **iii.** *p*_occ1_ of 0.7 and *p*_repl_ of 0.5. **iv.** *p*_occ1_ of 0.7 and *p*_repl_ of 0.3. **v.** *p*_occ1_ of 0.9 and *p*_repl_ of 0.9.

We proceeded by reducing the initial probability of the occupancy (*p*_occ1_) of *N* to 0.7 with a high probability of replenishment (*p*_repl_) between pulses of 0.9 (**Fig. 5Bi**) and a total *N* of 10 (occupied plus unoccupied) that remained constant among pulses as above (SES model). In this SES scenario, the first parabola reported the nominal value of the total *N* of 10. Notably, also in this scenario the points of the second pulse fell to the first parabola albeit at higher apparent *p*_N_ values. For both parabolas the reported *p*_N_ values were given by the nominal *p*_N_ values multiplied with the corresponding probabilities of occupancy of the *N*, i.e. *p*_N1_ = *p*_N_ * *p*_occ1_ and *p*_N2_ = *p*_N_ * *p*_occ2_. The occupancy during the second pulse was higher than during the first one due to the high probability of replenishment between pulses, which gave rise to the shift of the second points to apparently higher *p*_N_ values. The principle result that points of the first and the second pulse fall to the same parabola with their location being given by *p*_N_ * *p*_occ_ held for all different initial occupancies and different probabilities of replenishment tested (**Fig. 5Bii-v**). We can therefore conclude that changes in the occupancy of release sites are reminiscent of the changes in *p*_N_ simulated in the previous section (**Fig. 4D,E**) and are difficult to distinguish from them.

In summary (**Table 1**), in an MPFA with paired pulses the parabola will report the total *N* including those sites which are not occupied by a release ready SV. The location of the individual points along the parabola will depend on the product of the intrinsic *p*_N_ and the probability of occupancy. Importantly, points of the first and the second pulse fall to same parabola as long as the total *N* is constant. On the other hand, if the second parabola is wider than the first one, this is a strong indication for the recruitment of additional release sites. Finally, if the y-intercept from CumAna is larger than *N*_MPFA_ reported by MPFA this is a strong indication for the presence of a series-connected RP with finite size, whereas in the absence of the RP the y-intercept will be equal to *N*_MPFA_.

## Conclusions and Discussion

In this paper, we theoretically explored the insights that can be gained into the organization and dynamics of SV pools by combining the experimental methods CumAna and MPFA, including phenomena like overfilling and the recruitment of new release sites. We used computer simulations to guide our theoretical considerations and focused on two recent sequential vesicle pool models, the SOS (Doussau et al., 2017;Bornschein et al., 2019b) and the SES (Miki et al., 2016). We further covered a traditional PP model that does not harbor a series-connected RP (Wölfel et al., 2007;Mahfooz et al., 2016). In the following, we will summarize our main conclusions and discuss them in a broader context:

i. The presence of a series-connected RP but also parallel RRPs with SVs differing in their *p*_v_ give rise to a biphasic drop in quantal contents during sustained trains of APs with steady-state depression of synaptic responses. On the other hand, for a single RRP with direct replenishment from the quasi infinite reserve pool RSP this drop will be monoexponential, unless the replenishment rate gets reduced over time. Hence, the biphasic drop can indicate a more complex organization of SV pools but cannot distinguish between sequential and parallel arrangements.
ii. The y-intercept in CumAna reports the sum of RRP (=*N*_occ_) plus RP (SOS model: y(0) = RRP + RP) and would also include initially empty release sites (SES model: y(0) = RRP+ *N*_unocc_ + RP). In the PP model the y-intercept gives the total RRP (y(0) = Σ RRP_i_). If the RP would be added in the PP model, it would also show up in the y-intercept. The fit of the PP model with RP to experimental data could be superior to fits with the SOS or SES models. However, all these models typically have more parameters and equations than constrained by the experimental data and are therefore underdetermined already in their present forms.
iii. The first parabola of MPFA reports the total *N*, including initially empty sites, but excluding replenishment sites (*N*_MPFA_ = *N* \ RS). Hence, we suggest that if y(0) > *N*_MPFA_ this is a clear indication of the presence of a RP and RS, respectively. It should be noted that longer trains of APs are required. Briefer bursts (< 10 APs) may not exhaust the RP and the linear back extrapolation will yield a y-intercept that is closer to the estimate of *N* from the MPFA but has a strong dependency on the length of the burst. Experimentally y(0) > *N*_MPFA_ with long AP trains has been observed at different synapses (see below).
iv. If the parabola in a paired pulse experiment gets wider in the second pulse this results from a factual increase in *N* (see also Clements and Silver, 2000), whereas a mere increase in the occupancy of *N* does not change the shape of the parabola. Hence, MPFA with paired pulses provides a means to differentiate a factual increase in *N* from a mere increase in the occupancy of a fixed number of *N* as source of overfilling. Such increase in *N* has been observed experimentally for example at parallel fiber (PF) to Purkinje cell (PC) synapses (Valera et al., 2012;Brachtendorf et al., 2015), but not at PC – PC synapses (Bornschein et al., 2013).
v. For parallel pools, MPFA reports the total *N* summed over all sites (*N*_MPFA_ = Σ *N*_i_). Hence, for PP models y(0) will be equal to *N*_MPFA_, providing a means to distinguish the PP arrangement from the SOS or SES arrangements.

In summary, based on points (ii), (iii) and (v) we suggest that the y-intercept is greater than *N*_MPFA_ if presynaptic boutons harbor a RP and RS in series with the RRP (y(0) > *N*_MPFA_), while both are equal for parallel RRPs without RP (y_0_ = *N*_MPFA_). Biexponential decay of synaptic responses during AP trains thus indicates sequential or parallel pools (i), and the combined use of CumAna and MPFA offers the possibility to distinguish between SOS, SES on the one hand and PP on the other. Finally, MPFA with paired pulses provides a means to identify factual increase in *N* as opposed by a mere increase in the occupancy of *N*, i.e. it provides a means to distinguish between SOS and SES models (**Table 1**).

A recent variant of MPFA used trains of APs at ‘simple synapses’ that harbor only a single active zone, in combination with counting of individual release events by deconvolution (Miki et al., 2016;Miki, 2019). Variance mean analysis was extended there to cover also cumulative values, somewhat reminiscent of a combination of MPFA with CumAna. The quantification of the data relied on more complex modeling and provided evidence for sequential pools with incomplete initial release site occupancy (SES model). The aim of the present study was to probe if the classical and more simple versions of MPFA (Clements and Silver, 2000) and CumAna (Schneggenburger et al., 1999) can provide similarly deep insights into the organization of release sites and vesicle pools in the absence of complex modeling. Our results suggest, that a combination of the two methods with canonical parabolic or linear analysis can indeed provide detailed information about the organization of release at the active zone.

The results for the SOS and SES model will not be identical to another recent sequential model in which SVs reversibly shift their state from LS to TS before they can fuse (Neher and Brose, 2018;Neher, 2023). As in the SES model, empty sites (ES) occur in the LS/TS model and increasing occupancy of the TS is a major mechanism of facilitation. Similar to the RP in the SOS and SES models, SVs in LS will show up in the y-intercept, as the ES will do. However, different from vesicles in the RP, LS vesicles will also appear in the *N* of the MPFA. This means that in the LS/TS model y(0) and *N*_MPFA_ could be quite similar, just as in the PP model. Thus, while there are several similarities between the sequential models, there are also crucial differences between the SOS, SES and LS/TS models. In particular, vesicles of the RP or in LS are not identical (**Fig. 6**).

**Figure 6.**
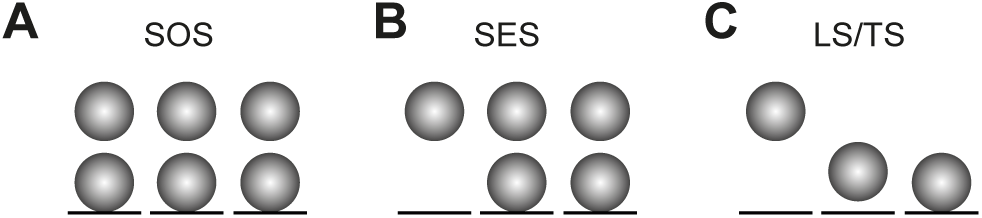
Three types of sequential models. **A.** In the SOS model all release sites (*N*, black lines) are occupied by SVs (spheres) at rest, forming the RRP (lower row). In addition there are replacement sites (RS) that are also occupied by SVs forming the RP (upper row). The reserve pool is omitted for clarity. During high-frequency synaptic activity *N* reversibly increases, thereby giving rise to facilitation (Dousseau et al., 2017). In the SOS model as shown here, the following parameters from CumAna and the first parabola of MPFA would be obtained: y(0) = 6, *N*_MPFA_ = 3. **B.** As in **A** but in the SES model not all *N* are occupied at rest. During high-frequency synaptic activity the occupancy of *N* but not the number of *N* itself reversibly increases, thereby giving rise to facilitation (Miki et al., 2016). In the SES model as shown here, the following parameters would be obtained: y(0) = 5, *N*_MPFA_ = 3. **C.** In the LS/TS model SVs reversibly shift between loosely (LS, middle) and tightly docked states (TS, right). There are also empty sites as in **B** (left). Fusion occurs only from the TS state. The LS/TS ratio is influenced by synaptic activity and determines short-term plasticity (Brose and Neher, 2018). Note that LS vesicles are attached to a release site while RP vesicles occupy replacement sites. Therefore, LS and RP vesicles are not identical. In the LS/TS model as shown here, the following parameters would be obtained: y(0) = 3, *N*_MPFA_ = 3.

In our simulations, released quanta were assumed to be perfectly synchronized, which is a simplification compared to real synaptic release as investigated in experiments. In the SOS and SES models it was further assumed that SVs from the RP do not contribute to the first release. In experiments CumAna and MPFA are typically based on the analysis of PSC amplitudes. Even with ultra-rapid recruitment (Miki et al., 2016;Doussau et al., 2017) in which SVs from the RP may even contribute to the first PSC it is more likely that these SVs contribute to the decay of the PSC rather than to its peak amplitude. In addition, in rapid-freezing EM studies the transient increase in the number of docked SVs was observed only after the stimulation (Kusick et al., 2022). Hence, we think that non-perfectly synchronized release will not affect the general conclusion of y(0) > *N*_MPFA_ for the SOS and SES models.

In the PP model also the low *p*_v_ pool got fully depleted in our simulations. Since the amplitude of the Ca^2+^ signal and the corresponding release rates rapidly drop with increasing distance from the Ca^2+^ channels (Bucurenciu et al., 2008;Bornschein et al., 2013;Schmidt et al., 2013;Nakamura et al., 2015;Bornschein et al., 2019b;Chen et al., 2024) this may not always be possible to achieve in experiments. In this case, the y-intercept will not report the sum of both RRPs but rather an intermediate value between the size of the high *p*_v_ pool and the sum of both pools, i.e. RRP_high *p*_ < y(0) < RRP_high *p*_ + RRP_low *p*_. Then, SVs from RRP_low *p*_ can contribute to the slope of the steady-state phase. In particular this may arise if *p*_v_ are strongly heterogeneous between SVs and/or if SV pools are very large, harboring thousands of SVs such as in the calyx of Held (Neher, 2015). Considering this possibility, the comparison of y- intercept and *N*_MPFA_ yields y(0) ≤ *N*_MPFA_ for the PP model. However, y(0) > *N*_MPFA_ as in the SOS and SES models will not occur.

Footprints of our conclusions can be found in experiments on different CNS synapses, including cerebellar, neocortical or brainstem synapses. Cerebellar granule cells (GCs) typically form only a single PF synaptic contact that harbors only a single active zone (Xu-Friedman et al., 2001) with their postsynaptic targets, including PCs and molecular layer interneurons (MLI). These ‘simple synapses’ are an intensively investigated model for a typical small cortical synapse (Pulido and Marty, 2017;Schmidt, 2019;Silva et al., 2021). MPFA in paired GC – PC recordings quantified *N*_MPFA_ and *p*_N_ to be ∼3 and 0.25, respectively (Schmidt et al., 2013). Strikingly, with this very limited immediate resource for transmitter release, PF synapses show long lasting high-frequency facilitation over up to 30 APs with initial paired pulse ratios of up to ∼3 (Valera et al., 2012;Brachtendorf et al., 2015;Doussau et al., 2017). MPFA with paired pulses at this synapse showed that the parabola in the second pulse got wider than in the first pulse, which was interpreted as a use-dependent factual increase in *N* as a major source of PPF at these synapses (Valera et al., 2012;Brachtendorf et al., 2015). Furthermore, the long lasting high-frequency facilitation of this synapse was explained by a sequential pool model with ultra-rapid, reversible increases in *N* (Doussau et al., 2017). The present considerations and simulations support this interpretation since they exclude that an increasing width of the parabola could result from a mere increase in the occupancy of a fixed number of release sites. However, they do not exclude that in addition to the increase in *N* there is also incomplete initial occupancy. Incomplete initial occupancy has been detected at MLI – MLI synapses (Trigo et al., 2012) and increasing occupancy of release sites during successive synaptic activations has been suggested at PF – MLI as mechanism of PPF, using the SES model (Miki et al., 2016). Since PF – PC and PF – MLI synapses are formed by the same presynaptic GC, it appears plausible that at PF terminals both mechanisms, increasing *N* and incomplete initial occupancy, are operational synergistically. For the interesting question of what constitutes an empty release site and more generally how the view of the release site has been changing in recent years, we refer to recent comprehensive reviews (Pulido and Marty, 2017;Neher and Brose, 2018;Silva et al., 2021;Kusick et al., 2022;Neher, 2023).

The prioritization of the sequential over the parallel model was not explicitly justified at PF synapses (but see (Doussau et al., 2017)). However, data from CumAna and MPFA show y(0) > *N*_MPFA_ at PF – PC synapses (Schmidt et al., 2013;Baur et al., 2015), which argues in favor of the sequential model. Similar findings come from an inhibitory cerebellar synapse (Chen et al., 2024) and a neocortical synapse formed between layer 5 pyramidal neurons (L5PNs (Bornschein et al., 2019a;Bornschein et al., 2019b). Interestingly, at the latter synapse evidence was found that the RP develops only during synaptic maturation between the first and third postnatal week in mice, thereby, changing the STP properties (Bornschein et al., 2019a).

The PP model was used to simulate the biphasic time course of release at another important model synapse, the calyx of Held (Wölfel et al., 2007). However, a biphasic time course of release can also arise from sequential pools of SVs (see point (i)) and it has recently been shown that the LS/TS model describes release and STP at the calyx very well (Lin et al., 2022).

Results on differences in STP are frequently considered to reflect differences in *p*_v_. The sequential pool models suggests an alternative interpretation (Neher, 2023). The dynamics of reversible vesicle priming, varying occupancy of release sites and the recruitment of new release sites make major contributions to STP. The simple theoretical framework proposed here provides a means to identify signatures of the RP without the need for complex computer simulations. It is based on the combination of two standard physiological methods, cumulative analysis of PSC amplitudes and multiple probability fluctuation analysis.

## Acknowledgment

We thank Stefan Hallermann for discussion of the manuscript. This work was supported by Deutsche Forschungsgemeinschaft grants to HS (SCHM1838/4-1, /6-1).

## Author contributions

HS designed research and performed simulations; GB, SB made the figures; HS, GB, SB wrote the manuscript.

## Abbreviations and glossary

|  |  |
| --- | --- |
| SV | Synaptic vesicle |
| PSC | Postsynaptic current amplitude; $PSC = p N q$ |
| QC | Quantal content quantifying the amount of release from the presynaptic terminal; $QC = p N$ |
| CumAna | Cumulative analysis, i.e. analysis of cumulative QCs or PSCs; a method for estimating the size of the RRP and the vesicular release probability |
| $y(0)$ | y-intercept of CumAna used to estimate the size of the RRP |
| MPFA | Multiple probability fluctuation analysis; a method to estimate the quantal synaptic parameters $p$, $N$ and $q$. |
| $p$ | Release probability |
| $p_v$ | $p$ per vesicle:<br>$p_v = PSC1/RRP$ or $QC1/RRP$<br>Calculated from CumAna as<br>$p_v = PSC1/y(0)$ or $QC1/y(0)$ |
| $p_N$ | $p$ per release site; reported by MPFA |
| $p_{occ}$ | Probability that a given release site is occupied |
| $p_{repl}$ | Probability that an emptied release site gets replenished between two pulses |
| $N$ | Binominal parameter reflecting the number of release sites including occupied and non-occupied sites |
| $N_{occ}$ | Release sites occupied by a release ready vesicle |
| $N_{unocc}$ | Unoccupied release site, empty release sites |
| $N_{MPFA}$ | $N$ reported by MPFA |
| $q$ | Quantal size: The postsynaptic response elicited by release of a single vesicle |
| $F$ | Synaptic failure rate summed over all release sites forming the total synaptic connection:<br>$F = (1 - p_N)^N$<br>$F = (1 - p_v)^{RRP}$ |
| $S$ | Synaptic success rate:<br>$S = 1 - F$ |
| $r$ | Rate constant of replenishment or recruitment |
| DS | Docking site; synonym of release site ($N$) |
| ES | Empty release site, i.e. a release site that harbors no SV |
| RS | Replacement site; SVs occupying these sites form the RP |
| LS | Loosely tethered state of a vesicle |
| TS | Tightly tethered state of a vesicle; SV in TS form the RRP in the narrower sense |
| RRP | Ready releasable pool of vesicles; equivalent to SVs in TS |
| RP | Exhaustible rapid replenishment pool formed by SVs that occupy recruitment sites; Ultra rapid transition of SVs from RP to RRP can cause overfilling and an increase in $N$; transition between RRP and RP is reversible on a slower timescale. Not identical to LS. |
| RSP, ∞ | Infinite reserve pool |

